# Nose-to-Brain Delivery of Cocaine in Mice: A Physiologically Based Pharmacokinetic Model Incorporating Olfactory Epithelial Transport, Mucociliary Clearance, and Blood-Brain Barrier Permeability

**DOI:** 10.64898/2026.11.12.766260

**Authors:** Maria F. Ramirez, Abhishek Gour, Emma K. Watson, Abhisheak Sharma, Daniel W. Wesson

## Abstract

Intranasal (IN) cocaine produces rapid central nervous system (CNS) effects despite generating substantially lower plasma concentrations than equivalent intravenous (IV) doses, suggesting a direct nose-to-brain pathway. The olfactory epithelium provides anatomical continuity for cocaine to pass between the nasal cavity and the brain. Quantifying the relative contribution of this pathway to total brain cocaine exposure requires a physiologically-based pharmacokinetic (PBPK) model that explicitly accounts for nasal mucosal architecture, epithelial transport barriers, and blood-brain barrier (BBB) permeability. Here we used tandem mass spectrometry to collect cocaine concentration-time data from the brain and plasma of awake mice administered IV or IN cocaine and leveraged that data to generate such a framework. Our results provide evidence that following IN cocaine, the olfactory pathway accounts for approximately half of the cocaine that reaches the brain. The mechanistic PBPK framework developed here — incorporating anatomically grounded nasal compartment separation, blood-flow-scaled mucosal absorption, tissue diffusion resistance, mucociliary clearance, plasma protein binding, and BBB permeability-surface area transport — substantially improves upon empirical one-compartment absorption models and produces parameter estimates with physiological interpretability. The proportionate efficiency of the olfactory route relative to systemic absorption may provide a pharmacokinetic basis for the rapid onset of IN cocaine’s CNS effects and has direct implications for understanding the psychoactive effects of cocaine.

**Significance statement:** Many drugs of abuse are inhaled, including cocaine, yet the mechanistic outcomes of inhaled cocaine are not understood compared to actions of cocaine following other ROAs. We established the pharmacokinetics of IN cocaine in awake mice, and developed a PBPK model to define the contributions of the olfactory pathway upon IN cocaine administration. Our findings demonstrate the contributions of a direct nose-to-brain pathway in mediating cocaine brain entry, thus, establishing the nasal ROA as a biological variable shaping pharmacokinetics and neurobiological outcomes of cocaine.

## Introduction

Cocaine is a highly addictive psychostimulant. The majority of cocaine users initiate drug-taking by insufflation of cocaine, which remains by many users, the preferred route of administration (ROA)^1,2^. Users of intranasal (IN) cocaine tend to snort it frequently and compulsively which holds high risk of overdose, as well as treatment complications^3–6^. The ROA dictates the pharmacokinetics of cocaine entry into the blood and the brain, which in turn highly influence the effects and physiological consequences of cocaine. Understanding the spatiotemporal dynamics of cocaine entry into the brain by means of the IN ROA is essential to elucidate the physiological and behavioral effects that are driven by cocaine and to develop mechanism-based therapeutics for cocaine use disorder, particularly for individuals who largely take cocaine IN.

The unique anatomical and physiological characteristics of the IN ROA dictate drug absorption and distribution. Unlike other ROAs, IN drugs can reach the CNS through two mechanisms: directly via sensory nerve pathways (*e*.*g*., the olfactory nerve or trigeminal nerve), or indirectly through systemic absorption and subsequent blood-brain barrier crossing. The nasal cavity’s rich vascularization, particularly in the respiratory epithelium, enables rapid absorption into systemic circulation^7–11^. The IN ROA also permits direct nose-to-brain transport that bypasses both systemic circulation and the blood-brain barrier^8,12–14^. Cocaine’s high lipophilicity allows it to be possibly transported through intracellular (neuronal uptake and axonal transport) and extracellular (perineural bulk flow along olfactory or trigeminal nerve pathways) mechanisms, allowing the drug to reach regions such as the olfactory bulb and brainstem^15–18^. The direct nose-to-brain transport, which bypasses systemic circulation entirely, may underlie the effects of insufflated cocaine.

The ROA-dependent kinetics of cocaine distribution and elimination in plasma following IN, oral, and intravenous (IV) cocaine have been established in humans^19–22^. These factors are less well-established in preclinical models, specifically in rodents. That said, in rats there is evidence for a direct nose-to-brain transport pathway, including that of significantly higher cocaine levels in the olfactory bulb compared to downstream brain regions following IN administration^13^. However, this prior work had important limitations: cocaine metabolism was not characterized, the animals were anesthetized, which likely influenced absorption kinetics, and the IN delivery method involved surgical isolation of the nasal cavity to prevent drainage to non-nasal regions, which restricted the ability to recapitulate the distribution of the IN ROA by hindering normal mucociliary clearance, drainage patterns, and potentially even blood flow to the region.

The ROA-specific absorption and distribution of IN cocaine also determines its metabolic profile^21,23^. Cocaine is primarily metabolized by hepatic and plasma esterases into two major inactive metabolites: benzoylecgonine and ecgonine methyl ester, which serve as canonical biomarkers of cocaine use^24,25^. The relative contributions of direct neural versus systemic pathways may influence the extent and timing of metabolite formation, further distinguishing IN from other ROAs.

Our goals for this study were threefold: first, to determine the ROA-specific distribution of cocaine by measuring peak plasma and brain concentrations across biologically relevant timepoints; second, to describe the metabolism of IN cocaine into its major metabolites; and third, to examine the regional distribution of cocaine across brain regions. To address these goals, we used a novel IN drug delivery device for mice recently developed by our group^26^ . This approach allowed us to deliver precise doses of IN cocaine in awake mice, without the limitations of the above noted study^13^. We compared IN cocaine profiles to those following IV administration, the most commonly used ROA in preclinical studies, to identify ROA-specific pharmacokinetic differences. Finally, we used the resulting data to develop a multicompartmental PBPK model in order to investigate nasal contributors to cocaine’s entry into the brain.

## Results

### Intranasal vs intravenous delivery of cocaine results in distinct pharmacokinetic profiles

Given that ROA critically influences cocaine pharmacokinetics and behavioral effects, we sought to determine the pharmacokinetics of IN cocaine administration—the most prevalent ROA in human drug users—and compare it to IV administration, the long-standing method used in preclinical studies. We delivered adult C57BL/6J mice an IN or an IV cocaine bolus of 8 mg/kg while they were awake and freely exploring their home cage. This dose was selected based upon our recent research which established this dose achieves robust behavioral outcomes in mice^27^. The infusion volumes and number of boluses were selected to ensure precise and comparable cocaine delivery across both ROAs (NAP^27^ and jugular catheter) and to maintain consistent delivery duration. Plasma and brain samples were collected at 1, 3, 6 or 12□min post-infusion for quantitative analysis of cocaine and its metabolites (benzoylecgonine and ecgonine methyl ester) using ultraperformance liquid chromatography–triple quadrupole mass spectrometry (UPLC–MS/MS) (**Fig 1A**; compound parameters for quantification of analytes using UPLC–MS/MS are represented in **Table 1**). Both plasma and brain samples from treated mice contributing data to all time points were included, as well as samples from control mice infused with physiological saline We found that cocaine distribution and pharmacokinetics markedly differed between the IN and IV ROA in both the plasma and brain (**Fig 1B**; three-way ANOVA, main effects of timepoint (*F*(4,48) = 16.94, *p* < 0.0001), ROA (*F*(1,48) = 107.6, *p* < 0.0001), and tissue (*F*(1,48) = 135.8, *p* < 0.0001). In the plasma, IV delivery produced rapid peak plasma cocaine levels at 1 min post-infusion (2432.7 ± 286.4 ng/mL), whereas IN delivery resulted in both a delayed and lower peak plasma level (323.1 ± 68.6 ng/mL) occurring at 3 min post-infusion. Similarly, brain cocaine levels reached a maximum concentration at 1 min following IV administration (7007.2 ± 1112.2 ng/g) while for IN administration maximum concentration was not observed until 6 min (1298.8 ± 247.2 ng/g), post-dose. Plasma and brain cocaine levels diverged significantly as early as 1 min following IV administration (**Fig 1B**, right panel); Tukey’s multiple comparisons IV plasma vs brain cocaine levels at 1 min *p* < 0.0001) but not until 3 min following IN administration (**Fig 1B**, left panel); Tukey’s multiple comparisons IN plasma vs brain cocaine levels at 3 min *p* < 0.0001), indicating that IV cocaine reaches the brain more rapidly than IN cocaine. Importantly, we found that the temporal dynamics of cocaine in brain vs. plasma were ROA-dependent, with IV and IN administration exhibiting distinct kinetic profiles that varied across timepoints (three-way ANOVA, significant interaction significant three-way interaction between timepoint, ROA, and tissue (*F*(4,48) = 6.46, *p* = 0.0003).

**Table 1.** Compound parameters for quantification of analytes using UPLC-MS/MS.

| Analytes | Parent (*m/z*) | Daughter (*m/z*) | Cone (V) | Collision (V) |
|---|---|---|---|---|
| Cocaine | 304.2 | 182.1 | 36 | 18 |
| Benzoylecgonine | 290.0 | 168.1 | 14 | 18 |
| Ecgonine methyl ester | 200.0 | 82.1 | 24 | 26 |
| Cocaine-*d*<sub>3</sub> (IS) | 307.1 | 185.1 | 24 | 18 |

**Figure 1.**
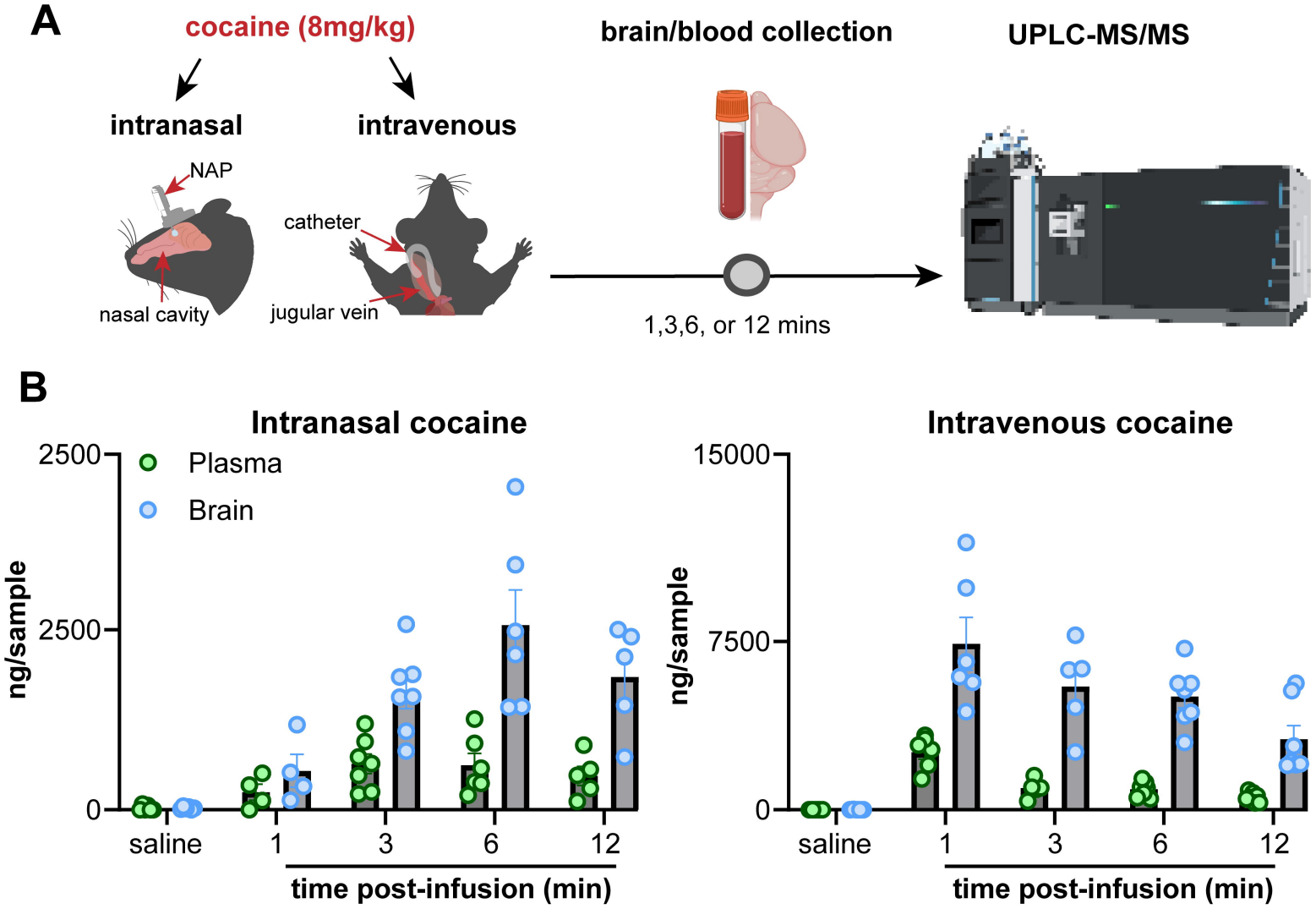
Intranasal and intravenous cocaine administration lead to distinct levels of cocaine in the brain and plasma. **A**. Schematic and timeline for collection of brain and plasma following intranasal (IN) vs. intravenous (IV) cocaine for chemical detection. Cocaine and metabolites were quantified using ultra-performance liquid chromatography-triple quadrupole mass spectroscopy (UPLC-MS/MS). Some schematic elements were created in BioRender. Wesson, D. (2025) https://BioRender.com/q73a907. **B**. Cocaine levels in plasma (ng/mL) and ipsilateral brain homogenates (ng/g) following IN (left) vs. IV (right) administration (8mg/kg). IV administration produced rapid peak levels in both plasma (2432.69 ± 286.40 ng/mL) and brain (7007.19 ± 1112.17 ng/g) at 1 min. IN administration resulted in delayed and lower peak levels in plasma (323.13 ± 68.58 ng/mL at 3 min) and brain (1298.81 ± 247.23 ng/g at 6 min). Three-way ANOVA revealed significant main effects of timepoint (*F*(4,48) = 16.94, *p* < 0.0001), route (*F*(1,48) = 107.6, *p* < 0.0001), and tissue (*F*(1,48) = 135.8, *p* < 0.0001), with significant interactions between timepoint × route (*F*(4,48) = 13.37, *p* < 0.0001), timepoint × tissue (*F*(4,48) = 9.50, *p* < 0.0001), route × tissue (*F*(1,48) = 73.32, *p* < 0.0001), and a three-way interaction (*F*(4,48) = 6.46, *p* = 0.0003); Tukey’s multiple comparisons showed differences between plasma vs. brain at 3 min following IN and at 1 min following IV administration (both *p* < 0.0001). Data presented as mean ± SEM, points = individual mice.

### Intranasal and intravenous delivery yield ROA-specific cocaine metabolite profiles

We next sought to understand the ROA-specific metabolism of cocaine by quantifying benzoylecgonine and ecgonine methyl ester, the primary metabolites of cocaine, in the same plasma and brain samples (**Fig 2**). In both IN and IV ROA, we observed that the levels of benzoylecgonine progressively increased over time in the plasma and brain following either ROA, resulting in higher levels in the plasma regardless of ROA (**Fig 2A**). Even though both ROA led to a similar increasing trend, the magnitude of benzoylecgonine levels in each tissue substantially differed between IV and IN. Mice that received IV cocaine resulted in higher levels of benzoylecgonine in both tissues. Differences in plasma levels occurred as early as 1 min (IV: 385.8 ± 34.7 vs IN: 19.5 ± 9.4 ng/mL), peaking at 12 min for both ROA (IV: 921.5 ± 65.7 vs. IN: 286.2 ± 66.1 ng/mL). On the other hand, both ROA yielded lower benzoylecgonine brain levels, differing between ROA at the 3 min timepoint (IV: 86.5 ± 12.4 vs IN: 16.7 ± 4.6 ng/g) and peaking at 3 min for IN (16.7 ± 4.6 ng/g) and 12 min for IV (153.2 ± 27.3 ng/g; benzoylecgonine; three-way ANOVA, significant three-way interaction between timepoint x ROA x tissue (*F*(4,48) = 6.37, *p* = 0.0003); Tukey’s multiple comparisons, IN vs IV brain at 3 min *p* = 0.0003, plasma at 1 min *p* < 0.0001). These findings are consistent with established cocaine pharmacokinetics, wherein benzoylecgonine represents the predominant metabolite and accumulates progressively over time, with higher plasma concentrations reflecting the primary site of hepatic metabolism^24,25^.

**Figure 2.**
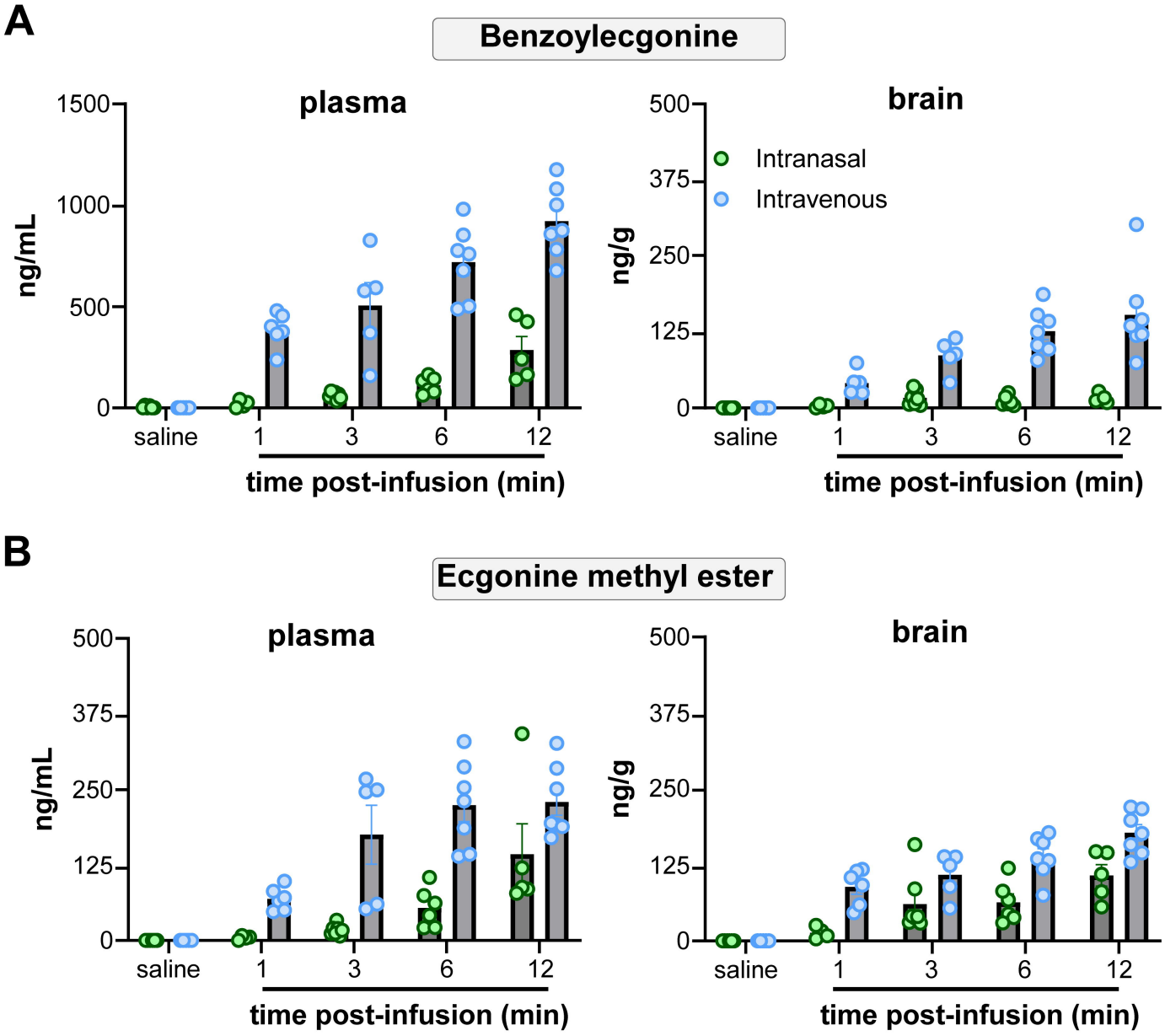
Intranasal and intravenous cocaine administration lead to different profiles of cocaine metabolites in the brain and plasma. Levels of **A**. benzoylecgonine and **B**. ecgonine methyl ester in plasma (ng/mL) and ipsilateral brain (ng/mg) following IN (green) vs. IV (blue) cocaine administration (8 mg/kg). Both metabolites showed progressive increases over time, with higher concentrations in plasma than brain regardless of route. IV administration consistently produced higher metabolite levels compared to IN. Benzoylecgonine levels differed between routes at 1 min in plasma (IV: 385.8 ± 34.7 vs. IN: 19.5 ± 9.4 ng/mL, *p* < 0.0001), with both routes resulting in peak levels at 12 min (IV: 921.5 ± 65.7 vs. IN: 286.2 ± 66.1 ng/mL), whereas in the brain, both routes showed differences at 3 min (IV: 86.5 ± 12.4 vs. IN: 16.7 ± 4.6 ng/g, *p* = 0.0003). Peak brain levels occurred at 3 min for IN (16.7 ± 4.6 ng/g) and 12 min for IV (153.2 ± 27.3 ng/g). Ecgonine methyl ester showed similar route-specific patterns, with significant differences emerging at 1 min in brain (IV: 89.1 ± 12.1 vs. IN: 13.4 ± 4.8 ng/g, *p* = 0.0007) and 3 min in plasma (IV: 505.6 ± 112.8 vs. IN: 62.1 ± 6.3 ng/mL, *p* < 0.0001), peaking at 12 min for both plasma and brain (IV: 921.5 ± 65.7 vs. IN: 286.2 ± 66.1 ng/mL) and brain (IV: 178 ± 13.4 vs. IN: 107.8 ± 17.8 ng/g). Three-way ANOVAs revealed significant main effects and interactions for both metabolites (benzoylecgonine: timepoint *F*(4,48) = 50.43, route *F*(1,48) = 216.2, tissue *F*(1,48) = 198.2, three-way interaction *F*(4,48) = 6.37; ecgonine methyl ester: timepoint *F*(4,48) = 42.80, route *F*(1,48) = 146.9, tissue *F*(1,48) = 214.8, three-way interaction *F*(4,48) = 11.15; all *p* < 0.0001 except benzoylecgonine three-way *p* = 0.0003). Data presented as mean ± SEM, points = individual mice, same mice as in Fig 1.

Concurrently, we analyzed ecgonine methyl ester in our samples, another primary metabolite of cocaine formed through esterase-mediated metabolism in liver and plasma^24,25^. We found an increasing trend in the levels of ecgonine methyl ester in both ROAs (**Fig 2B**). Similarly to benzoylecgonine, metabolism of cocaine into ecgonine methyl ester in the plasma and the brain was specific to each cocaine ROA. We found that levels of ecgonine methyl ester at each timepoint were higher in the plasma than in the brain for both ROAs. Levels of ecgonine methyl ester significantly differed between IN and IV in the brain as early as 1 min (IV: 89.1 ± 12.1 vs IN: 13.4 ± 4.8 ng/g), and in the plasma at the 3 min time point (IV: 505.6 ± 112.8 vs IN: 62.1 ± 6.3 ng/mL). Peak ecgonine methyl ester occurred at 12 min in the plasma (IV: 921.5 ± 65.7 ng/mL vs IN: 286.2 ± 66.1 ng/mL) and the in brain (IV: 178 ± 13.4 ng/g vs. IN: 107.8 ± 17.8 ng/g) for both ROAs (Ecgonine methyl ester; three-way ANOVA, interaction between timepoint x ROA x tissue (*F*(4,48) = 11.15, *p* < 0.0001); Tukey’s multiple comparisons, IN vs IV brain 1min *p* = 0.0007, plasma 3min *p <* 0.0001). Collectively these results indicate that cocaine metabolism exhibits distinct kinetics depending on its’ ROA, with IV delivery consistently producing higher metabolite levels across all timepoints and tissues examined.

### Brain distribution

We also examined cocaine and cocaine metabolite levels in the following regions of interest within the brain (ROIs): the olfactory bulb, due to its proximity to the nasal cavity and potential indicator for direct nose-to-brain transport; the ventral striatum and midbrain, as these regions are central to the mesolimbic reward system mediating many of cocaine’s psychoactive and behavioral effects^28–30^; and the cerebellum, as an auxiliary region in the most distal hindbrain for comparison of cocaine distribution patterns. We examined across different timepoints the relative distribution of cocaine in those ROIs by comparing the detected levels of cocaine over the total amount of cocaine in the contralateral brain (**Fig 3**). We observed striking differences in the relative accumulation of cocaine when delivered IN vs IV, related to the kinetics of how cocaine enters the brain when delivered IN vs IV. When delivered IV we observed a more uniform distribution of relative cocaine levels in the ROIs across timepoints. At peak IV brain levels (1 min timepoint), the relative cocaine distribution was 19.5% ± 4.6 in the olfactory bulb, 28.6% ± 1.7 in the ventral striatum, 24.6 % ± 3.6 in the midbrain and 27.7% ± 0.9 in the cerebellum (**Fig 3A**, right panel). In contrast, following IN cocaine infusions, we found a more compartmentalized distribution of cocaine in the different ROIs across the timepoints. Over 30% of cocaine across all time points was localized in the olfactory bulb, and it reached its highest level at the 3 min timepoint (37.6% ± 9.3), while in the ventral striatum was (16.6% ± 3.4), midbrain (23.9% ± 3.6), and cerebellum (23.6% ± 5.1) (**Fig 3A**, left panel).

**Figure 3.**
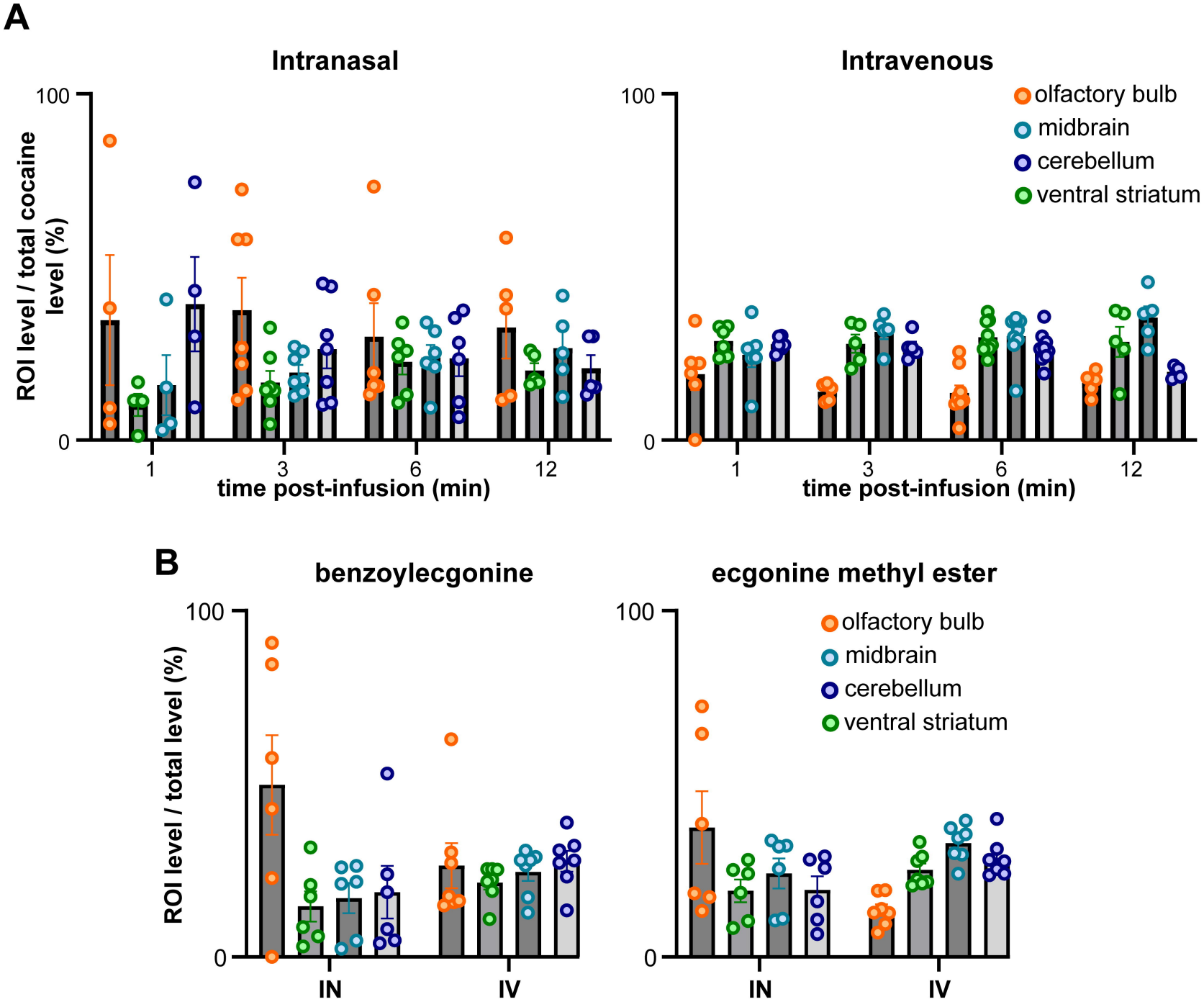
Intranasal and intravenous cocaine administration result in region-specific differences in cocaine and metabolite bioavailability in the brain. **A**. Relative distribution of cocaine across brain regions of interest (ROIs) in the contralateral hemisphere at the different timepoints following intranasal (IN) (left) or intravenous (IV) (right) administration. Values are expressed as the percentage of total compound levels within the contralateral hemisphere (ROI level /total contralateral hemisphere level). Following IV administration, peak brain levels of cocaine (1min timepoint), showed relatively uniform distribution across regions (19.5% ± 4.6 in the olfactory bulb, 28.6% ± 1.7 in the ventral striatum, 24.6 % ± 3.6 in the midbrain and 27.7% ± 0.9 in the cerebellum). In contrast, IN administration resulted in preferential accumulation in the olfactory bulb (>30% across all timepoints, peak 37.6% ± 9.3 at 3 min) with lower levels in the ventral striatum (16.6% ± 3.4), midbrain (23.9% ± 3.6), and cerebellum (23.6% ± 5.1). **B**. Relative distribution of benzoylecgonine (left) and ecgonine methyl ester (right) in the contralateral brain at 6 min (peak ipsilateral cocaine levels). Two-way ANOVA revealed a significant ROI × route interaction for ecgonine methyl ester (*F*(1.312,14.43) = 5.16, *p* = 0.031) and a trend for benzoylecgonine (*F*(1.5,19.8) = 3.126, *p* = 0.07). The most pronounced differences occurred in the olfactory bulb: IN administration resulted in substantially higher metabolite accumulation (29.9 ± 10% benzoylecgonine; 37 ± 10% ecgonine methyl ester) compared to IV (13.6 ± 2%; 13.4 ± 2%, respectively; *p* = 0.0039 for ecgonine methyl ester).

Since cocaine levels peaked at 6 min following IN delivery (compared to 1 min for IV), we examined whether ROA-dependent differences in cocaine distribution also influenced regional brain metabolism patterns. We compared cocaine metabolite levels across brain regions and found that the distribution of benzoylecgonine and ecgonine methyl ester differed between ROA, with the most pronounced differences observed in the olfactory bulb (**Fig 3B**). While benzoylecgonine showed a trend toward ROA-dependent regional differences (**Fig 3B**, left panel), two-way ANOVA, ROI × ROA interaction: *F*(1.5,19.8) = 3.126, *p* = 0.07), ecgonine methyl ester exhibited significant ROA-dependent regional distribution (**Fig 3B**, right panel), two-way ANOVA, ROI × ROA interaction: *F*(1.31,14.43) = 5.16, *p* = 0.031). IV administration resulted in 13.6 ± 2% of benzoylecgonine and 13.4 ± 2% of ecgonine methyl ester in the olfactory bulb, (*p* = 0.0039 for ecgonine methyl ester). Whereas, IN administration resulted in preferential metabolite accumulation in the olfactory bulb, with 29.9 ± 10% of benzoylecgonine and 37 ± 10% of ecgonine methyl ester localized to this region. These findings suggest that IN delivery promotes greater metabolite formation in the olfactory bulb relative to other brain regions, while IV delivery produces more uniform metabolite distribution.

### Modelling the contributions of the nasal pathway to cocaine’s entry into the brain

Finally, we used the above concentration-time data of cocaine to generate a compartmental model to investigate the contributions of the nasal pathway to cocaine’s entry into the brain. Since each mouse contributed one timepoint, in this classic sparse cross-sectional design, we treated the data as population PK. We leveraged prior nasal models to embed key anatomical and cytoarchitectural constrains about the mouse nasal cavity^31–33^ and also physiochemical constraints about cocaine, in-part informed by the MS-MS results presented above. The anatomical and cytoarchitectural constrains included subdivisions of the nasal airway including their thicknesses, serial regions in the nasal cavity, tissue compartments (mucus, epithelial, subepithelial), and fluid dynamics that are known to be altered by breathing rates. These constraints also include known differences in metabolic functions of both the respiratory and olfactory mucosa. The model (see Methods for details) was developed as a two-stage model – first generating simulation data based upon the real IV data, and second generating simulation data based upon the real IN data.

As shown in **Figure 4**, the two-stage PBPK model adequately described the observed/real plasma and brain cocaine concentration-time profiles. For IV administration, we observed a decline in plasma cocaine concentrations, with a steep drop between 1 and 3 mins suggesting rapid distribution into brain and peripheral tissue, followed by a shallower elimination phase through 12 mins. Cocaine concentrations in the brain were consistently 3–5-fold higher than plasma following IV delivery at all time points, consistent with the parameterized brain partition coefficient and *K*_*p*,brain_ =1.769 *and f*_u_ × *K* _*p*,brain_ =7.0 ^34^ with the fixed permeability–surface area product (*PS*_brain_ =1.0 mL/min/mL) capturing rapid equilibration across the mouse BBB (**Fig 4A**).

**Figure 4.**
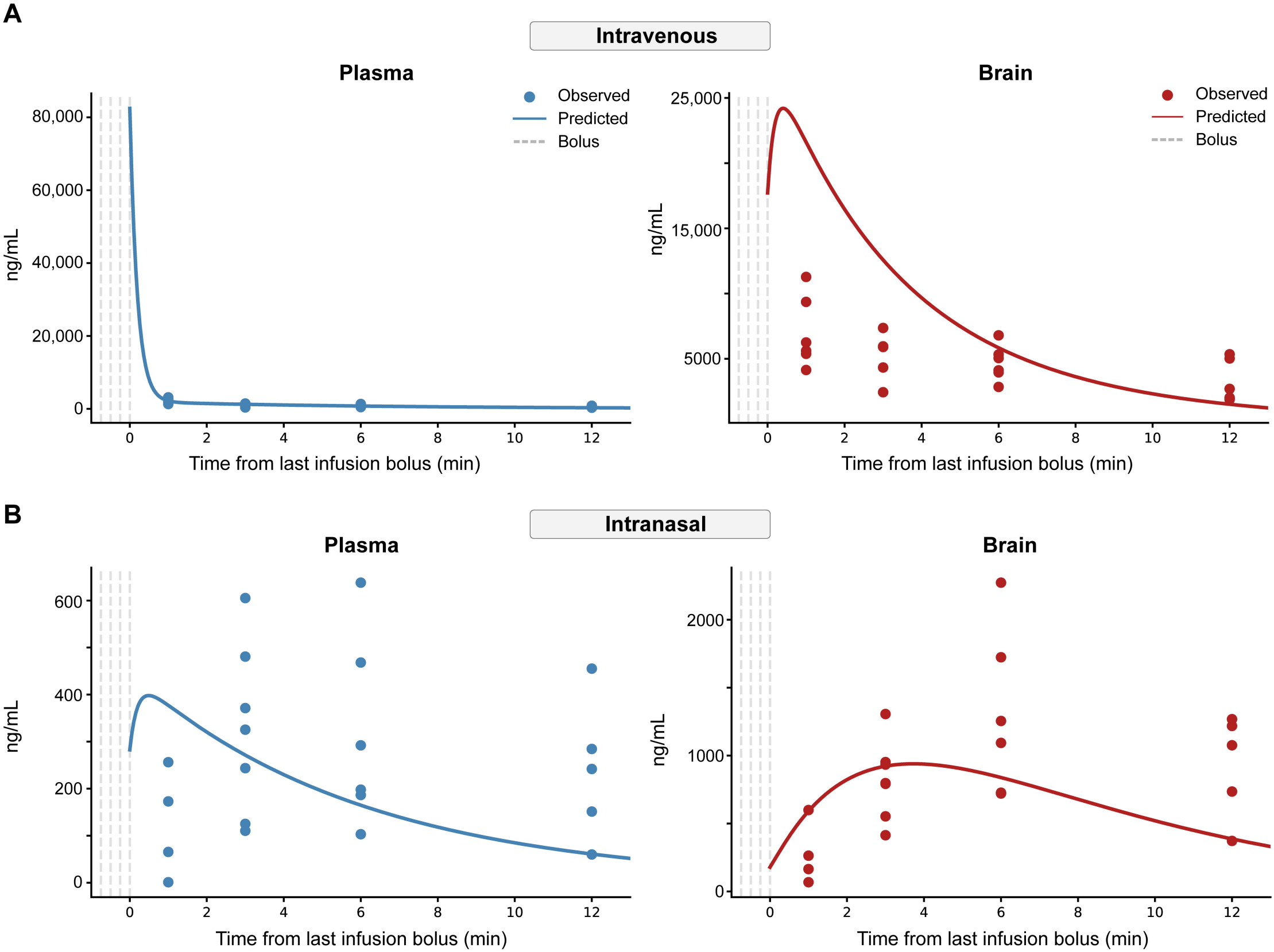
Physiologically-based pharmacokinetic (PBPK) model of IN vs IV cocaine. Two-stage PBPK model predictions (lines) compared to observed/real (dots) plasma (blue) and brain (red) cocaine concentrations in mice following IV and IN administration of identical nominal doses (dashed lines indicate delivery time of each cocaine bolus). **A**. IV ROA model produced a biphasic plasma profile (left panel) with rapid initial distribution over 1-3 mins into brain and peripheral tissues, followed by slower elimination phase through 12 minutes. Brain concentrations (right panel) were consistently 3-5-fold higher than plasma across all timepoints. B. IN ROA model based on anatomical and cytoarchitectural constraints of the nasal cavity indicates plasma concentrations (left panel) 10–20-fold lower than IV, reflecting slow nasal absorption (*ka_resp_eff* ≈ 0.0169 min□^1^) and substantial mucociliary clearance. Brain concentrations (right panel) after IN dosing were disproportionately elevated relative to plasma, with progressively increasing brain-to-plasma ratios from 1-12 minutes, indicative of direct olfactory delivery.

The second stage of the model captured IN dynamics. Following IN administration, we observed plasma cocaine concentrations 10–20-fold lower than IV at equivalent time points despite identical nominal doses, suggesting slow nasal absorption (*ka_resp_eff* ≈ 0.0169 min□^1^) and substantial mucociliary loss that we estimated at approximately 92.2% loss of the respiratory epithelium deposited dose and a 97.5% loss of the olfactory epithelium deposited dose (**Fig 4B**, left). By partitioning the nasal cavity into anatomically distinct respiratory and olfactory compartments and governed by vascular perfusion fractions (see Methods), we found that the model reproduced the slowly rising IN plasma profile in a manner that a single nasal compartment alone could not achieve.

We observed that brain concentrations following IN administration were disproportionately elevated relative to plasma, with brain-to-plasma ratios increasing progressively from 1 to 12 mins — a temporal pattern incompatible with a purely systemic absorption model and consistent with sustained direct olfactory delivery. By incorporating an cocaine directly into the brain volume (*v*_3_), the model captured this rising ratio across all four olfactory neuroepithelial compartment that bypasses systemic circulation and the BBB to deliver to the anatomical ceiling of *F*_olf_ =0.55 (reflecting the ∼50% olfactory surface coverage of the observation times (**Fig 4B**, right). With initial deposition on the olfactory epithelium constrained rodent nasal cavity) fitting Stage 2 yielded a baseline absorption rate constant *k*_*a*,base_ of 0.148± 0.042 min^-1^and a direct olfactory transfer rate constant k_olf_of 8.09±4.48Jx10^-4^ min^-1^.

By applying the tissue diffusion correction factor of 1/7 derived from Bush et al. ^31^ — reflecting the approximately 7-fold greater viscosity of cytoplasm relative to water — we found that the effective respiratory absorption rate resulted in systemic absorption rate constants of

*k* _*a*,resp_eff_ =0.0169 min^-1^ from respiratory mucosa and k_*a*,olf_sys_ =0.0042 min^-1^ from olfactory submucosa. Mucociliary clearance (k_mcc_ =0.20min^-1^) competed directly with mucosal absorption resulting in an estimated 92.2% first-pass loss of the respiratory-deposited dose to the (*e*.*g*., to the nasopharynx) prior to systemic entry. Within the olfactory compartment, competing mucociliary clearance and submucosal absorption limited the fraction of the total administered dose delivered directly to the brain to approximately 0.22%. We reason this influence of *k_mcc* to mean that mucociliary clearance further reduces the contribution of the systemic route to cocaine entry into the brain.

Finally, we iteratively excluded olfactory versus peripheral pathways (*Cb_olfactory* vs *Cb_systemic*) and compared those to total brain cocaine (*Cb_total*) through a pathway decomposition analysis (**Fig 5A**). We used area under the curve (AUC) to examine concentration-time data and synthetically expanded the time series from 12 to 30 minutes to capture prolonged resolution of the contributions of each nose-versus peripheral-pathways to cocaine levels detected in the brain. From this we can estimate that ∼50% of cocaine detected in the brain of a mouse arises from the nose-to-brain pathway (**Fig 5B**). Comparing the full model to a systemic-only model (*k*_olf_ =0) resulted in a modest improvement in fit (reduced *X*^*2*^ improved from 1.51 to 1.42; AUC from 19.1 to 17.4). Overall, the PBPK model supports the prediction that despite extensive mucociliary clearance, direct olfactory transport provides a meaningful contribution to early and sustained direct nose-to-brain transport of cocaine upon IN exposure.

**Figure 5.**
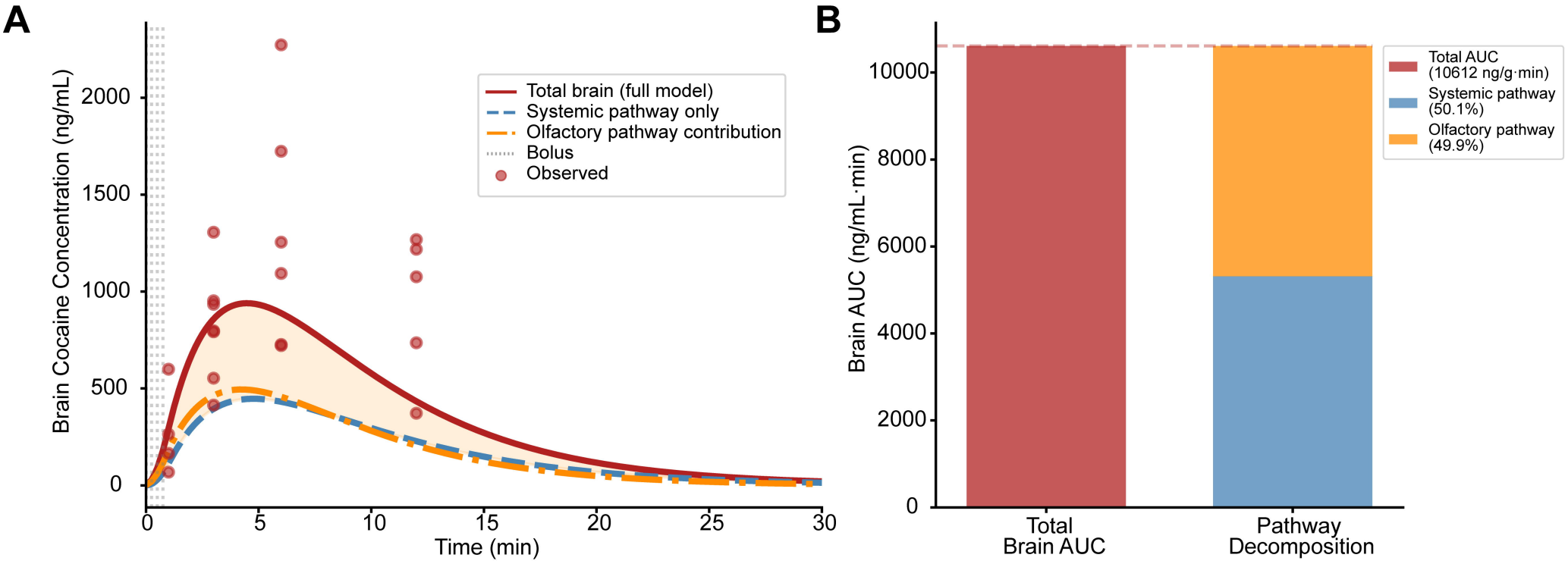
PBPK model pathway decomposition demonstrates predominant nose-to-brain transport of cocaine following intranasal administration. **A**. Area under the curve (AUC) following iterative exclusions of olfactory versus peripheral pathways demonstrated the contributions of each pathway to brain cocaine levels. Dashed lines indicate delivery time of each cocaine bolus. **B**. Pathway decomposition outcomes indicating that from the total AUC = 10612 ng/g·min, approximately 49.9% of cocaine detected in the brain following IN delivery originates through the olfactory pathway.

## Discussion

The ROA fundamentally shapes the pharmacokinetic profiles and the neurobiological consequences of cocaine^35–38^. The nasal cavity has privileged anatomical access to the brain^39^ and a rich enzymatic environment capable of local metabolism^40^, yet we do not fully understand the temporal dynamics, brain and blood distribution, or the metabolic kinetics that the nasal ROA confers to cocaine as it enters the body. Here, by focusing on early-time points upon delivery, we establish the comprehensive pharmacokinetic profile of IN cocaine in awake mice, and from those data, develop a PBPK model that we used to investigate the putative mechanisms whereby IN cocaine reaches the brain.

### ROA-specific temporal dynamics of cocaine elevations

We used a high-resolution temporal sampling strategy with early timepoints at 1, 3, 6, and 12 mins post-infusion. Previous rodent studies used extended sampling intervals (10-20 min) to uncover ROA-specific temporal dynamics – reporting peak cocaine levels occurring around 10 min for IN and nearly immediately for IV administration^13,41^. Here we uncovered similar rapid peak plasma and brain cocaine levels following IV delivery. Through the more rapid sampling strategy herein, we found that IN delivery resulted in cocaine plasma levels occurring within 3 mins, which while slower than following IV cocaine, is more rapid than indicated through prior reports. The PBPK model recapitulated these distinct kinetic profiles (**Fig 4**), confirming that the delayed plasma concentrations following IN administration result from slow nasal absorption and substantial mucociliary clearance upon cocaine uptake.

We found higher levels of cocaine in the brain than in the plasma for both routes, consistent with prior IN/IV comparisons of brain to plasma ratios in rats^13^, and as reported following IP delivery in mice^41^. The higher brain levels could be explained by the lipophilicity of cocaine, which allows it to enter the brain rapidly and, subsequently, to concentrate in the brain’s lipid compartments^41^. The PBPK model (**Figs 4 & 5**) also showed this relationship, confirming that IV brain concentrations remain consistently 3-5-fold higher than plasma due to rapid blood-brain barrier equilibration. For the IN ROA, we observed brain-to-plasma ratios that increased progressively over time—a pattern that could only be captured by incorporating a dedicated olfactory compartment that bypasses both systemic circulation and the BBB. This rising brain-to-plasma ratio across all four timepoints indicates that IN cocaine transport occurs in-part by sustained direct olfactory delivery and not purely by systemic distribution.

Users report euphoria within 3–5 minutes of nasal cocaine use^20,37^ and preclinical models show behavioral and neural effects occurring also within that timeframe^27,42^. Yet here we found peak plasma concentrations following IN cocaine are delayed. This suggests that the rapid psychoactive and behavioral effects of cocaine may be driven by the rate of increase in brain cocaine concentrations rather than peak plasma levels, or potentially by direct nose-to-brain transport, which could deliver cocaine to reward-relevant brain regions prior to systemic distribution. The PBPK model provided mechanistic insights into ROA-dependent pharmacokinetic differences and may help resolve the apparent temporal disconnect between plasma levels and behavioral effects. The model successfully captured, albeit with differences between real and simulated outcomes in some aspects, the distinct kinetic profiles we observed. The model revealed that the elevated brain concentrations following IN administration cannot be explained by systemic absorption alone. Instead, the model indicates that olfactory pathway input accounted for approximately half of brain cocaine exposure over 30 minutes, yet that only 0.22% of the total dose reaches the brain directly from nose-to-brain pathway, bypassing systemic circulation entirely (**Fig 4**). It is important to note that our model was created with the kinetic assumption that cocaine is only passively diffused, but we acknowledge that cocaine can undergo a tandem carrier-mediated transport process which robustly influences BBB permeability^43^ and may influence this interpretation. Additionally, our model does not compartmentalize systemic peripheral tissue by organ-specific absorption rates, which may affect predicted systemic brain delivery. It is possible incorporating carrier-mediated BBB kinetics and organ-resolved peripheral compartments would refine the model’s accuracy and, more importantly, would clarify the robustness of the direct pathway estimate.

### ROA dictates distribution of cocaine across brain regions

To elucidate how IN cocaine enters and is distributed across the brain, we examined the relative distribution of cocaine across brain regions of interest (ROIs). We observed ROA-dependent differences in the relative distribution of cocaine related to the kinetics of brain entry (**Fig 3A**). IV administration resulted in a uniform distribution of cocaine across ROIs at its peak level. This uniform distribution pattern is consistent with systemic delivery where cocaine enters the brain primarily via the bloodstream and distributes somewhat evenly across vascularized regions.

In contrast, IN cocaine administration produced a compartmentalized distribution pattern with preferential accumulation in the olfactory bulb compared to either the ventral striatum, midbrain, or cerebellum. This compartmentalized distribution was particularly pronounced at early timepoints, suggesting that IN delivery provides direct access to olfactory structures that bypasses initial systemic circulation. The large accumulation of cocaine in the olfactory bulb would suggest nose-to-brain transport direct into the olfactory bulb. The accumulation of cocaine in the cerebellum could be indicative of transport along trigeminal nerve pathways. The trigeminal nerve extensively innervates the nasal mucosa and projects to brainstem nuclei that have connections to the cerebellum, potentially facilitating drug distribution to this region, allowing the drug to reach such regions within minutes^15–18^. Through the PBPK model **(Fig 5)**, and by iteratively excluding olfactory versus peripheral pathways, we estimated that direct olfactory input may account for half of brain cocaine exposure. This finding supports our regional distribution results, confirming that the preferential olfactory bulb accumulation and compartmentalized distribution patterns indicate a prominent role for direct olfactory delivery, versus that largely through systemic circulation, in mediating IN cocaine’s brain entry.

The IN region of interest data had notable across-animal variability not apparent in the IN hemi-brain data. This is not unexpected given the small volume of the samples collected and this variability might be further explained by the state of the animal. The respiratory state of the animal at the time of administration, whether engaged in active sniffing or quiescent breathing— could significantly influence drug deposition patterns within the nasal cavity^40^. Active sniffing increases airflow velocity and turbulence, potentially enhancing drug deposition in the olfactory region and promoting direct nose-to-brain transport, whereas quiescent breathing may favor deposition in the respiratory region with subsequent absorption via mucosal blood.

### Metabolism of cocaine is dependent on ROA

Cocaine metabolism also differed by ROA. Both IN and IV administration produced progressively rising benzoylecgonine and ecgonine methyl ester in plasma and brain (**Fig 2**), with benzoylecgonine as the dominant metabolite in both routes, as expected from established cocaine pharmacokinetics^44,45^. The nasal mucosa is rich in metabolic enzymes, including carboxylesterases responsible for cocaine hydrolysis to both benzoylecgonine and ecgonine methyl ester^40^, and studies in bovine nasal tissue have demonstrated benzoylecgonine formation in both olfactory and respiratory epithelium. A notable finding from our results was the ROA-dependent regional distribution of metabolites within brain regions (**Fig 3B**). Following IN administration, both benzoylecgonine and ecgonine methyl ester were most elevated in the olfactory bulb, whereas IV administration resulted in more uniform distribution across brain regions. This regional specificity is particularly notable given that benzoylecgonine has limited blood-brain barrier permeability, thus, limited access to the brain from systemic circulation^34^, suggesting an alternative source of cocaine metabolite formation.

The different peak times for brain metabolite levels between ROAs in the brain (3 min for IN benzoylecgonine vs. 12 min for IV) further support the involvement of direct transport pathways. The earlier peak following IN administration suggests local formation and faster access to the olfactory bulb, rather than delivery via systemic circulation following hepatic metabolism. In contrast, the later, higher peak following IV administration reflects the time required for systemic distribution, hepatic metabolism, and subsequent brain delivery from systemic circulation. Further, while traditional pharmacokinetic models assume exclusive plasma and organ-based metabolism of cocaine. The present regional distribution data herein suggest that nasal mucosal metabolism may represent an additional, anatomically-specific source of metabolite formation that contributes directly to brain exposure following IN administration.

### Conclusions

Through a combination of IN cocaine delivery and quantification, and computational approaches, here we establish a comprehensive pharmacokinetic profile of IN cocaine in awake mice. Our work here in mice aligns with prior studies human studies demonstrating the fundamental principle that IN cocaine absorption is slower than when delivered IV^19–22^, and follows a compartmentalized distribution, particularly pronounced at early timepoints in regions such as the olfactory bulb, suggesting that IN delivery provides, in part, direct access to olfactory structures that bypasses initial systemic circulation. We uncovered evidence that the ROA itself is a impactful biological variable shaping both pharmacokinetics and likely neurobiological outcomes. Through the PBPK model, we demonstrate a prominent role for direct olfactory delivery, versus that largely through systemic circulation, in mediating IN cocaine’s brain entry. Overall, these findings elucidate the pharmacokinetic mechanisms underlying cocaine’s rapid CNS actions, closely recapitulating the route of administration used by humans, and providing insights into the biological determinants of its abuse potential.

## Methods

### Animals

Adult C57BL/6J mice (n=65, 38 males, 27 females, 2-6 months of age) originating from breeder stock at the Jackson labs (strain #000664; RRID:IMSR_JAX:000664, Bar Harbor, ME), were bred and maintained at the University of Florida. A subset of the mice (n=24, 15 males, 9 females) contributed data towards analysis of IN cocaine and its metabolites in a prior study^26^. Mice were housed in a temperature-controlled vivarium, on a 12:12hr light-dark cycle with *ad libitum* access to food and water. All experimental procedures were approved by the University of Florida Institutional Animal Care and Use Committee.

From the total sample size, a total of 4 mice (3 male, 1 female) that received IN cocaine, and 1 male mouse that received IV cocaine, where identified as outlier outliers by GraphPad PRISM’s (version 10) Robust Regression and Outlier Removal (ROUT) method (Q=1%) and were excluded from analyses. Further, from the group that received IN cocaine, 1 female was excluded due to plasma cocaine levels being negligible than those of other samples in the same group, while hemisected brain concentrations remained high. Similarly, 1 male mouse was excluded from analyses due to brain cocaine and metabolite concentrations being negligible than those of other samples at the same time point. For both of those mice, hemisected brain concentrations remained high, comparable to other hemisected brains in the same group, and consistent with the other hemisphere used for dissection of regions of interest.

### Surgical Procedures

For all surgical procedures, mice were anesthetized with 2-4% Isoflurane in oxygen (IsoFlo®, Patterson Veterinary, Houston, TX) and mounted on a stereotaxic frame while body temperature was maintained on a 38°C heating pad underneath their body. Meloxicam (20mg/kg, Patterson Veterinary) was subcutaneously (s.c.) administered followed by lidocaine (lidocaine, 3mg/kg, s.c., Patterson Veterinary, Houston, TX) to the scalp/midscapular region/upper chest area before incision.

### Implantation of nasal access port (NAP) for intranasal delivery

Mice were implanted with the NAP as we previously described^26^. A 1mm diameter craniotomy was drilled through the right nasal bone, positioned 1mm anterior to the frontal/nasal fissure and 0.5mm lateral. The NAP was lowered through the craniotomy, and secured with dental cement (Teets cold cure, Diamond Springs, CA). A removable aluminum cap with a magnetic insert was placed over the implant to protect it while not in use.

### In-dwelling jugular catheter implantation for intravenous delivery

Mice were implanted with chronic in-dwelling jugular catheters (Instech labs, Plymouth Meeting, PA). Two incisions were made: a small (<1 cm) incision in the midscapular region along the midline created a subcutaneous pocket for catheter passage, while a second incision (<1 cm) extended diagonally from the upper right chest to the neck. The right jugular vein was localized by blunt dissection and ligated at the superior end. The vein was partially incised with spring scissors, and a sterile 25-gauge polyurethane catheter (Instech Labs) was advanced toward the heart. The catheter was secured with suture, the incisions were closed with wound clips, and a removable aluminum cap was placed over the infusion port, which was attached to the catheter and extruded from the skin over the midscapular region.

Upon completion of either surgery the mice were single housed in a clean cage with a heating pad to aid recovery. Mice were given Meloxicam (20 mg/kg, s.c., Patterson Veterinary) for a 3-day post-operative period following the surgery.

### Patency testing

Patency of NAPs was established the day prior to cocaine infusion. Mice were briefly anesthetized under 2-4% Isoflurane in oxygen and 0.1 mL of saline was infused through the NAP to ensure a small amount of fluid could travel through the NAP and into the nasal cavity as evidenced by a visible saline solution drop exiting the ipsilateral nostril.

Jugular catheter patency was maintained daily from the day of surgical implantation. Mice were carefully restrained and syringe was attached to the catheter port to flush the catheter with 0.05 mL of 0.33 mg/kg heparin/gentamicin solution. Following brain and plasma detection (see below), a methylene blue solution was injected into the port for a visual confirmation of patency.

### Cocaine delivery

Cocaine hydrochloride was generously provided by the Drug Discovery Program at the National Institute on Drug Abuse and was dissolved ^26^ in sterile physiological saline (0.9% NaCl). Concentration was calculated w/v and adjusted to achieve desired concentration per infusion based on body weight of the animal (mg/kg/inf). For delivery of cocaine, mice were tethered via the NAP or jugular catheter to a female mating connector attached to polyurethane (PU) tubing (0.43□mm inner diameter × 0.94□mm outer diameter, 25 gauge) line. The line was filled with cocaine and attached to a syringe pump (R-210, Razel Scientific Instruments, Saint Albans, VT) which was controlled to deliver a single bolus of 8 mg/kg of cocaine (IN: 1 µL, X4, every 15s, IV: 6 µL X4, every 15s). Mice were placed in a chamber filled with isoflurane vapor for tissue and plasma collection over ice at 1, 3, 6, or 12 min, post-infusion.

### Ultra-performance liquid chromatography-triple quadrupole mass spectrometer (UPLC-MS/MS)

Tissue and plasma samples were prepared for cocaine detection, and ran for UPLC-MS/MS as we recently described in detail^27^. Blood was collected via cardiac puncture into EDTA tubes containing pre-aliquoted potassium fluoride (2% w/v final concentration; Thermo Fisher Scientific). The brain was subsequently perfused with potassium fluoride via cardiac injection and both brain and plasma stored at -80°C. Samples were analyzed against freshly prepared calibration and quality control standards in respective matrix following the US-FDA guideline for bioanalytical method validation^46^. These samples were processed by addition of 80μL of acetonitrile (Thermo Fisher Scientific) containing internal standard (cocaine-*d*_3_; IS, 2 ng/mL, Millipore Sigma St. Louis, MO). Samples were mixed, filtered, centrifuged at 1500 rpm, 4°C for 5 min, and injected onto Waters Acquity UPLC coupled with Xevo® TQ-S micro triple quadruple mass spectrometer. Chromatographic separation was achieved using a Waters Acquity UPLC BEH C18 column with a VanGuard pre-column of a similar chemistry (Waters Co, Milford, MA). The optimized linear gradient program for quantitation of cocaine and its metabolites similarly followed our prior work^26^. Quantification of cocaine and its metabolites was performed using TargetLynx software. Compound parameters for quantification of analytes using UPLC-MS/MS are displayed in **Table 1**.

### Multicompartment model of IN cocaine kinetics

We developed a two-stage PBPK model to describe the time course of cocaine concentrations in plasma and brain. We informed the overall model architecture using established nasal dosimetry PBPK frameworks for the rat and mouse nasal cavity^31,47,48^ and we structured the model as separate submodels for each route of administration sharing a common systemic pharmacokinetic core as follows:

### IV submodel

We constructed the IV submodel as three compartments arranged in series: plasma (V1), peripheral tissue (V2), and brain (V3). Following IV administration, we instantaneously added the entire dose/bolus to the plasma compartment at each scheduled bolus time, calculating the initial plasma concentration immediately following each bolus as:

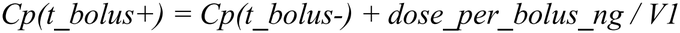

where V1 is the individual animal’s plasma volume scaled linearly for a 25g reference mouse (V1_ref = plasma volume ∼1.8mL blood × 0.47 plasma fraction ≈ 0.85mL). We similarly scaled hemi-brain volume V3 and peripheral tissue volume V2 (V3_ref = 0.250 mL, V2_ref = 1.5 mL) in proportion to individual body weight relative to the 25g reference ^30,51^

Because tissue collection occurred at 1, 3, 6, and 12min following the completion of the dosing regimen, we translated observational sample times to the internal model clock (*t*_model_ =*t*_obs_ +0.75 min) to preserve kinetic continuity from the first administered bolus.

We calculated drug transfer between compartments using a system of ordinary differential equations (ODEs) formulated in concentration units (ng/mL):

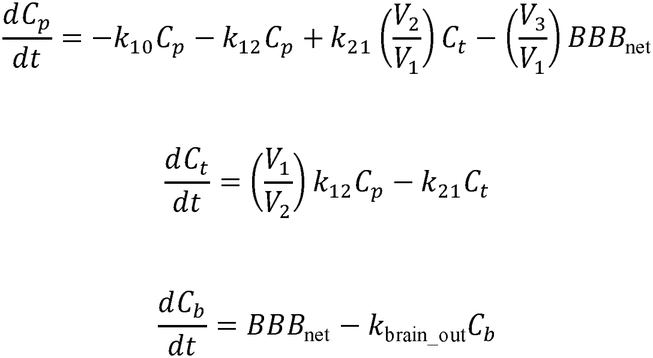

where *k*_10_ is the first-order central systemic elimination rate constant, and *k*_1_ and *k*_12_ govern bidirectional distribution between plasma and peripheral tissues. We applied volume ratios (*v*_2_/*v*_1_ and *v*_1_/*v*_2_) to distribution terms to maintain strict conservation of mass between compartments of differing volume.

We described transport across the BBB mechanistically using a permeability-surface area product (*PS_brain* = 1 mL/min per mL brain) to represent passive diffusion capacity across the BBB endothelium, rather than using an empirical rate constant. We formulated bidirectional BBB flux as:

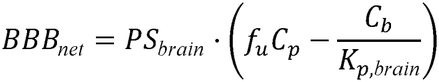

Additionally, *f*_*u*_ represents the unbound fraction of cocaine in mouse plasma (*f*_*u*_ =0.65)^52^. We parameterized the equilibrium partition coefficient as *K* _*p*,brain_ 7.0/*f*_*u*_ ≈ 10.77, consistent with the stable ∼7-fold brain-to-plasma concentration ratio reported across 5 to 60min following systemic cocaine administration in mice^34^. We captured residual brain clearance via a term *kbrain_out* representing non-BBB-mediated elimination of cocaine from the brain, including bulk cerebrospinal fluid drainage and local cerebral metabolism.

### IN submodel

We extended the IV systemic core with an anatomically structured nasal absorption module comprising two distinct epithelial compartments to construct the IN submodel, informed by the mucosal architecture of the rodent nasal cavity as described by Andersen et al.^32^ and the CFD-PBPK framework of Bush et al ^31^.

### Nasal compartment architecture

To maintain mass balance without relying on unmeasured nasal tissue subcompartment volumes, we tracked cocaine mass within the nasal cavity as discrete amounts (ng): an anterior respiratory mucosal amount (A_resp_) and a dorsal/posterior olfactory mucosal amount (A_olf_). At each of the four sequential administrations (t = 0, 0.25, 0.5, and 0.75min), parameter F_olf_, representing the fraction of the applied liquid bolus initially contacting olfactory the administered bolus mass was partitioned between the two mucosal surfaces according to the neuroepithelium:

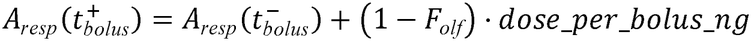

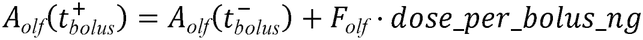

We constrained F_*olf*_ to an upper boundary of 0.55, consistent with the approximately 50% anatomical surface coverage of olfactory epithelium reported for rodents^47^.

### Nasal absorption and clearance kinetics

Drug cleared from the nasal epithelial surfaces via competing first-order absorption and mucociliary transport pathways:

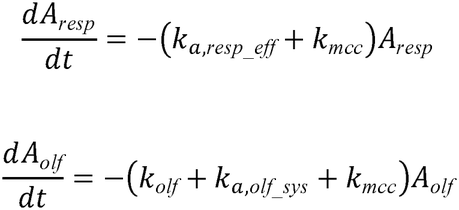

Systemic absorption rate constants from the respiratory mucosa (k _*a*,*resp_eff*_) and olfactory submucosa (k_*a*,*olf_sys*_) were defined from a fitted baseline absorption rate *(k*_*a* ,*base*_) scaled by regional vascular perfusion fractions *(F*_blood_resp_ =0.80, *F*_*blood_olf*_ =0.20)^31^ and an empirical tissue diffusion correction factor *(D*_*tissue_factor*_ 1/7). We derived the tissue diffusion correction factor of 1/7 from the validation studies of Bush et al., who demonstrated that diffusion coefficients in the epithelial and subepithelial layers. We attributed this reduction to the approximately 6.9-fold greater viscosity of cytoplasm relative to water^31^, consistent with the Stokes-Einstein relationship in which diffusion coefficient is inversely proportional to solvent viscosity. We note that Bush et al. considered this correction as the single most influential parameter in their model. In contrast, Bush et al demonstrated that gas-phase mass transfer coefficients and blood flow distribution had negligible effects on overall uptake, which led to our decision to fix these parameters at literature values rather than treat them as free fitting parameters.

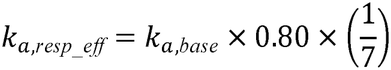

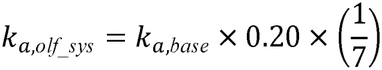

We defined mucociliary clearance as the physical transport of the mucus blanket and any dissolved or suspended material posteriorly toward the nasopharynx, where it is swallowed rather than absorbed nasally^30^. We set a rate constant of *k_mcc* = 0.20 min□^1^, representing a central estimate for rodent mucociliary transport rates, and we held this term fixed since we considered it a physiological property of the nasal epithelium independent of cocaine pharmacokinetics. We acknowledge that respiratory epithelium, with its dense population of ciliated cells, is the primary site of mucociliary clearance activity, while olfactory epithelium has sparser ciliation^32^ , however, we applied a uniform *k_mcc* to both compartments as a conservative simplification.

### Olfactory direct-to-brain pathway

Drug deposited on the olfactory neuroepithelium entered the modeled brain compartment directly via perineural routes bypassing systemic circulation and the BBB:

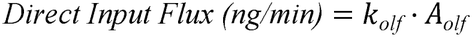

where *k*_*olf*_ represents the first-order direct transfer rate constant. This direct mass entry was converted to brain concentration per unit time by dividing by the modeled receiving brain volume *(v*_3_):

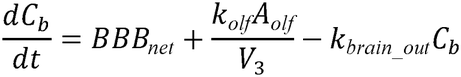

Systemic venous inputs from both mucosal sites entered the central plasma compartment scaled by plasma volume *(v*_1_):

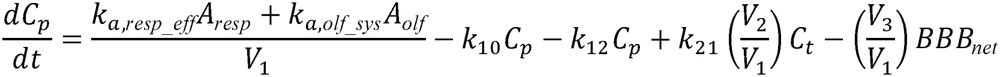

We based the biological rationale for this pathway on the anatomical continuity between olfactory sensory neurons, whose dendrites project into the nasal lumen (upon the nasal turbinates) and whose axons project through the cribriform plate directly into the olfactory bulb, providing perineural spaces through which lipophilic drugs such as cocaine can travel by convective and diffusive mechanisms. While we recognize that olfactory tissue carboxylesterase activity documented by Bogdanffy and Taylor^33^ — with *Vmax* values for olfactory tissue ranging from 89 to 165 µmol/min/mg protein, 2–6-fold higher than respiratory tissue — represents a competing metabolic loss term that would reduce effective olfactory delivery, we did not explicitly model this pathway in the current analysis and we address it as a limitation below.

### Parameter estimation

We conducted parameter estimation sequentially using a two-stage framework in Python (lmfit)^49^. Residuals were evaluated on a logarithmic scale:

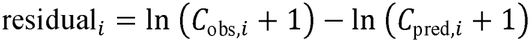

Observed brain tissue homogenate concentrations (*c*_*b*_, reported in ng/g wet tissue) were converted to volumetric tissue units (1g wet brain ≈ 1mL tissue)^50^, ensuring dimensional congruence with modeled *c*_*b*_ (ng/mL)

We employed a two-stage sequential fitting procedure to estimate pharmacokinetic parameters from the experimental ‘real’ data. In Stage 1, we estimated *k*_10_ and *k*_brain_out_ exclusively from IV plasma and brain profiles (*n* =50 observations across 25 mice), while fixing distribution constants (k_12_= 1.00 min^-1^, *k*_21_

=0.20 min^-1^) and BBB permeability (*PS*_brain_ 1.0 mL/min/mL) to maintain parameter identifiability under sparse population sampling. In Stage 2, we fitted the baseline nasal absorption rate constant *k*_*a*,base_ and direct transport constant *k*_olf_ against IN plasma and brain profiles (*n*=44 observations across 22 mice), with *F*_olf_ fixed at its anatomical upper bound of 0.55, while holding the systemic shared parameters fixed at their Stage 1 values, thereby preventing the more variable IN data from distorting the core pharmacokinetic structure.

For optimization, we utilized global differential evolution followed by Levenberg– relative and absolute tolerances of 10^-6^and 10^-9^, respectively. Marquardt local refinement. Systems of ODEs were solved using the LSODA algorithm^51^ with

### Pathway AUC decomposition

To quantify the fraction of total brain exposure resulting from direct olfactory transport versus systemic BBB permeation, we conducted an in silico source-labeling simulation across a 30-min window post-dosing. The full model (*AUC*_total_) was simulated with all pathways active. Systemic-only brain exposure (*AUC*_systemic_) was computed by setting direct transfer to zero (*k*_olf_ · *A*_olf_ =0) while preserving identical nasal mucosal depletion and systemic absorption. Direct olfactory exposure was derived by subtraction (*AUC*_olfactory_ *AUC*_total_ − *AUC*_systemic_) and was verified by confirming that decomposition discrepancy remained < 0.01% across all integrated using the composite trapezoidal rule (np.trapezoid). Exact mass-balance consistency integration time steps.

### Statistical Analysis

All data were processed using semi-automated steps whenever possible, including processing data from mixed treatment groups at the same time. Statistical comparisons were analyzed in GraphPad Prism which included repeated measures (rm) and/or two-way and three-way ANOVAs followed by Tukey’s multiple comparison corrections when needed. All data are reported as mean ± SEM unless noted otherwise.

## List of nonstandard abbreviations

IN: intranasal
IV: intravenous
PBPK: physiologically-based pharmacokinetic
ROA: route of administration
UPLC–MS/MS: ultraperformance liquid chromatography–triple quadrupole mass spectrometry

## Data availability

The data that support the findings of this study are available on request from the corresponding author. The data are not publicly available due to privacy or ethical restrictions.

## Code availability

The PBPK model can be accessed here: [https://colab.research.google.com/drive/1P8qL1r7mHqa-g41P0e7oxBphoC8jQxim?usp=sharing]

## Acknowledgements

We thank Ellyse Thomas for expert animal care assistance. This work was supported by R01DA049449, R21DA063991, and funds from the University of Florida to D.W.W.. M.F.R is supported by T3DC2015994 and F31DA066224.

## Author Contributions

Conceptualization: MFR & DWW

Methodology: MFR, AG, AS, DWW

Investigation: MFR, EW, AG, DWW

Visualization: MFR & DWW

Funding acquisition: MFR & DWW

Supervision: DWW

Writing – original draft: MFR & DWW

Writing – review & editing: MFR, EW, AG, AS, DWW

## Declaration of generative AI and AI-assisted technologies in the manuscript preparation process

During the preparation of this work, the authors used Anthropic’s Claude for troubleshooting of the PBPK model code for use in Python. The authors reviewed and edited the output as needed and take full responsibility for the content of the published article.

